## Supplementary figures for "Multiple molecular events underlie stochastic switching between two heritable cell states in a eukaryotic system"

### Table S1

**White to Opaque switching**

| Genotype | Plates | Colonies | Sectors | Percent Switch | Fold Change |
|---|---|---|---|---|---|
| WT | 11 (3,3,5) | 1340 | 17 | 1.27 (+/- 0.306) | - |
| Wor1 motif deletion | 11 (3,3,5) | 1790 | 8 | 0.447 (+/- 0.158) | 2.83 down |
| Wor1-GFP | 11 (3,3,5) | 948 | 491 | 51.8 (+/- 1.62) | 40.8 up |
| Wor1-mGFP | 11 (3,3,5) | 1603 | 31 | 1.93 (+/- 0.344) | 1.52 up |

**Opaque to White switching**

| Genotype | Plates | Colonies | Sectors | Percent Switch | Fold Change |
|---|---|---|---|---|---|
| WT | 11 (3,3,5) | 1926 | 87 | 4.52 (+/- 0.473) | - |
| Wor1 motif deletion | 11 (3,3,5) | 1159 | 224 | 19.3 (+/- 1.16) | 4.28 up |
| Wor1-GFP | 11 (3,3,5) | 1633 | 0 | <0.06 | >75 down |
| Wor1-mGFP | 11 (3,3,5) | 1707 | 52 | 3.05 (+/- 0.42) | 1.48 down |

Table S1 – Switching frequencies
Switching frequencies as calculated by counting sectors on colonies grown on solid ager plates. Data comes from three independent experiments containing multiple plates per strain. Numbers of colonies and sectors are pooled across plates. The percent of colonies with at least one sector is calculated. Standard error is the standard error of a sample proportion ($\sqrt{[p (1-p) / n]}$).

### Figure S1

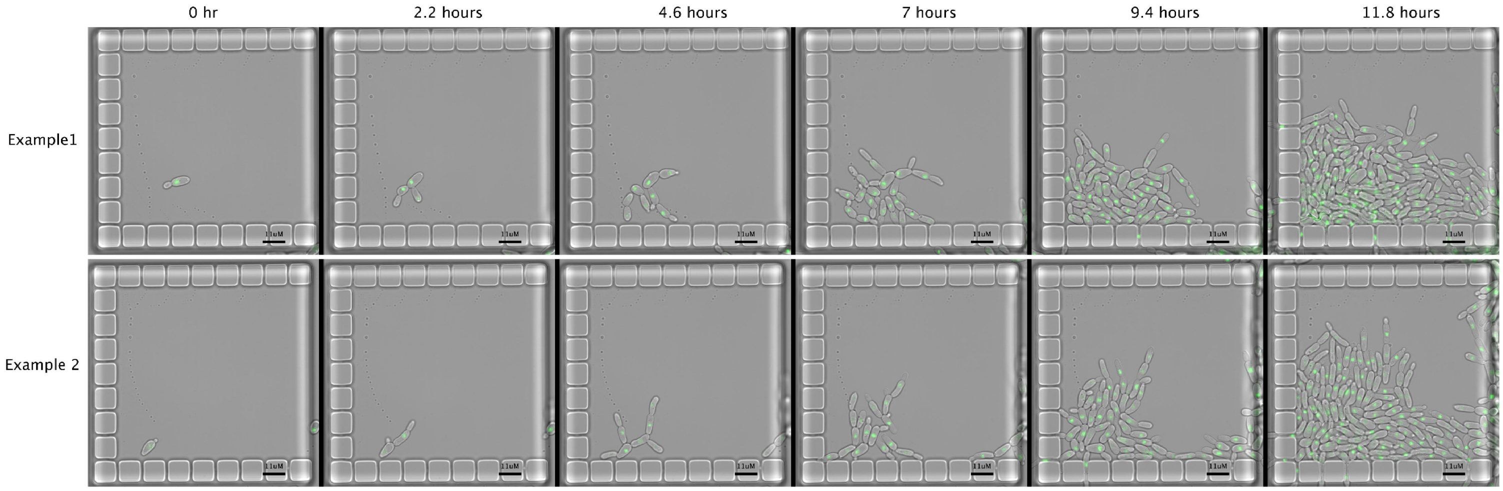


Figure S1 – Opaque cell growth in microfluidic traps
Two examples of microfluidic traps with growing opaque cells. Strain contains Wor1-GFP fusion protein.

### Figure S2

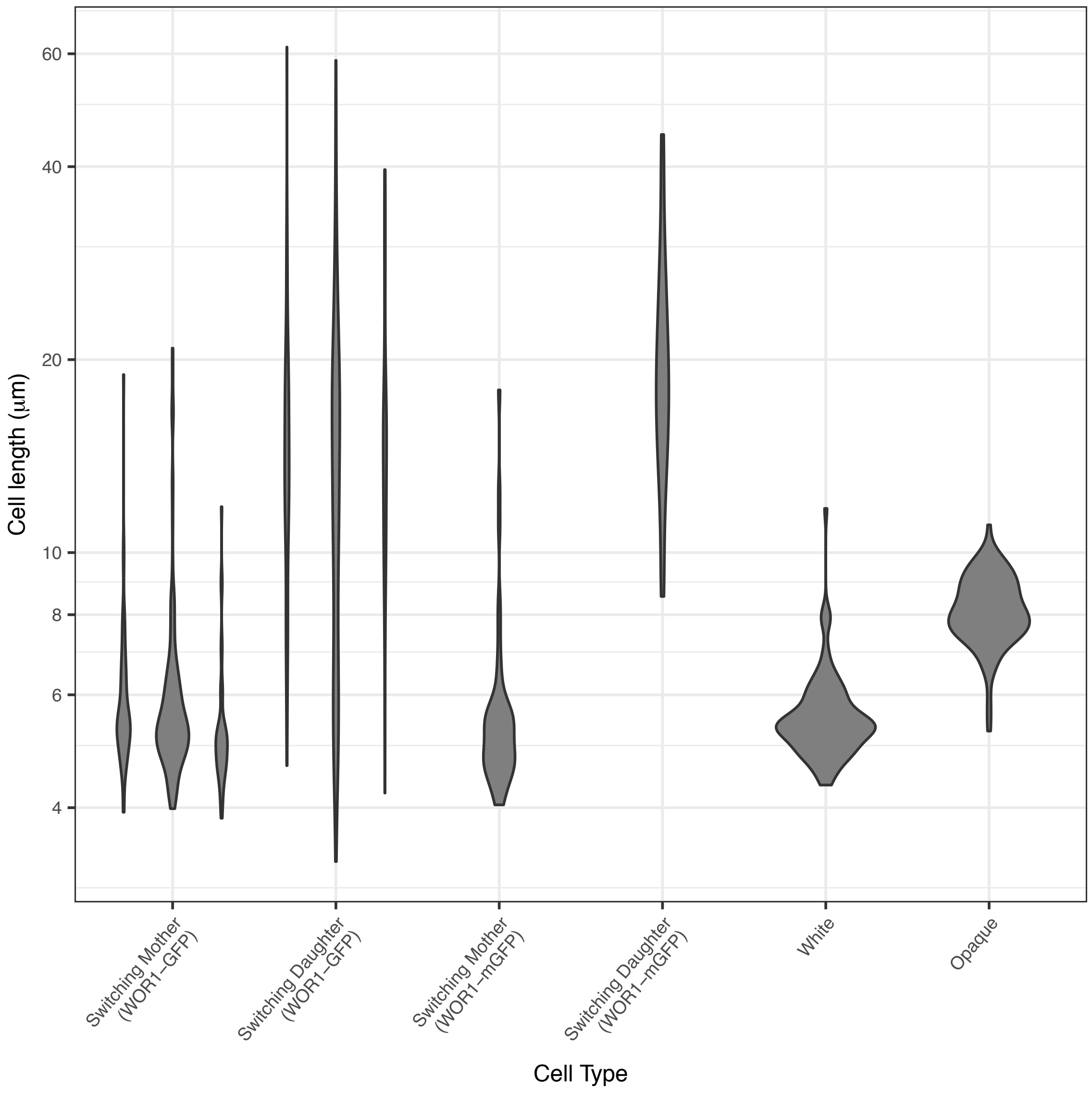


Figure S2 – Cell length distributions
Cell length distributions (violin plots) of different cell types, including switching cells containing Wor1-mGFP. For the Wor1-GFP strain, data comes from three independent experiments that were pooled in Figure 2.

### Figure S3

A

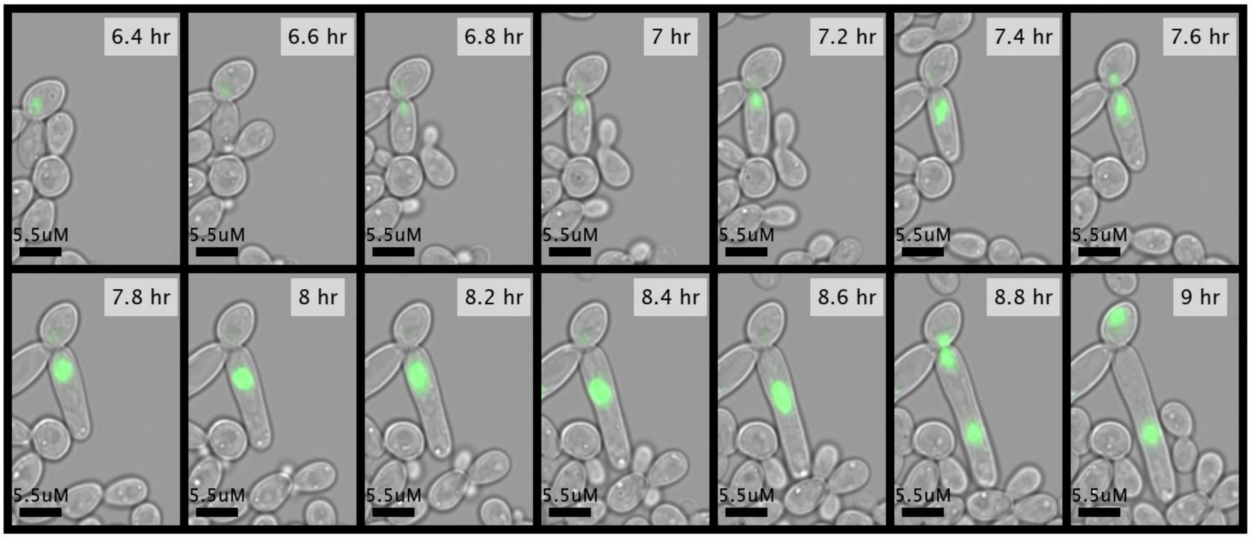


B

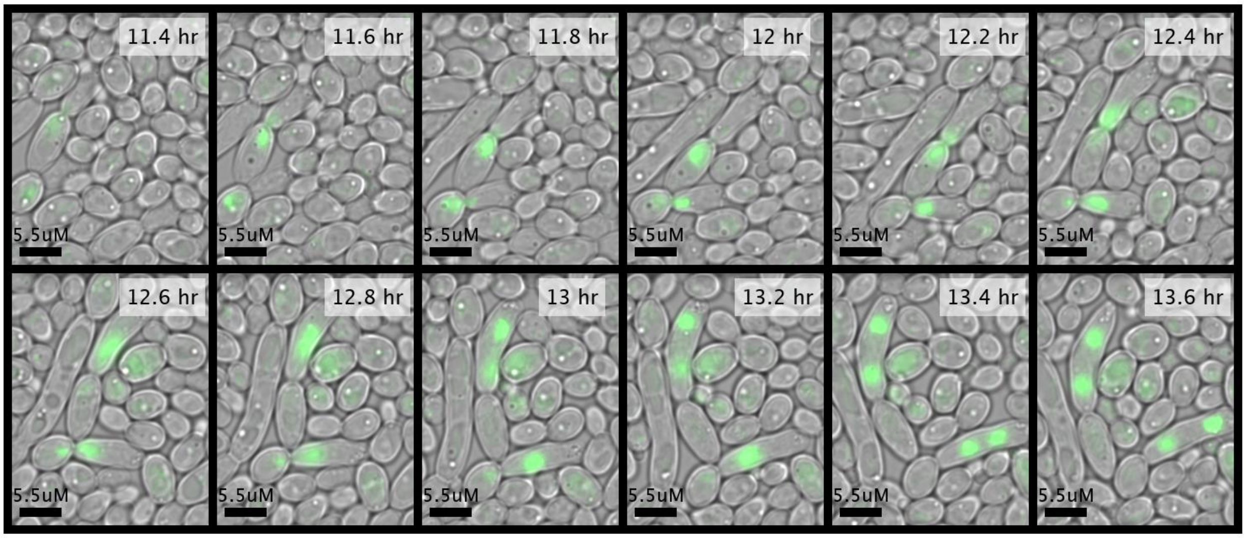


Figure S3 – Nucleus loss during mitosis in switching cells
A) Representative example of a cell division where the mother cell nucleus enters the elongated daughter cell but returns to the mother cell. Strain contains Wor1-GFP fusion protein. B) Representative example of cell divisions where the mother cell nucleus enters the elongated daughter cell and remains in the daughter cell, creating a polyploid cell. There are two cell divsions with this pattern in this example. Strain contains Wor1-GFP fusion protein.

### Figure S4

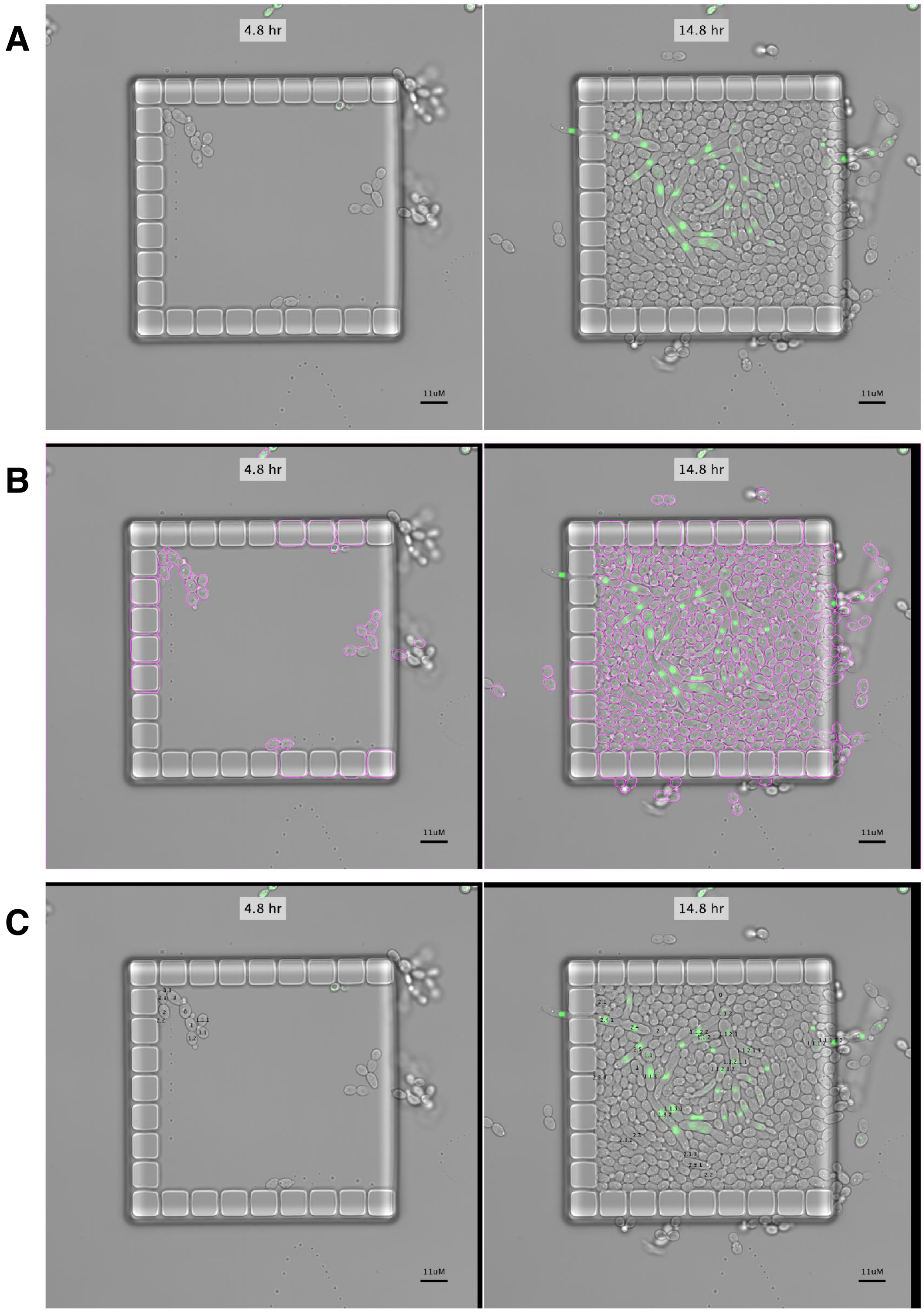


Figure S4 – Semi-automated image analysis pipeline

A) Representative original images of two time-points for one field. B) Same images as (A) after adjustment for (x,y) movement (note black regions at image edge) and automated segmentation of cells (magenta colored lines defining cell borders). C) Same images as (A) and (B) with overlay of manually defined cell identities, pedigree information is explicitly encoded in cell names (e.g., cell “2.1” is the first daughter of cell “2” which in turn is the second daughter of cell “0”).

### Figure S5

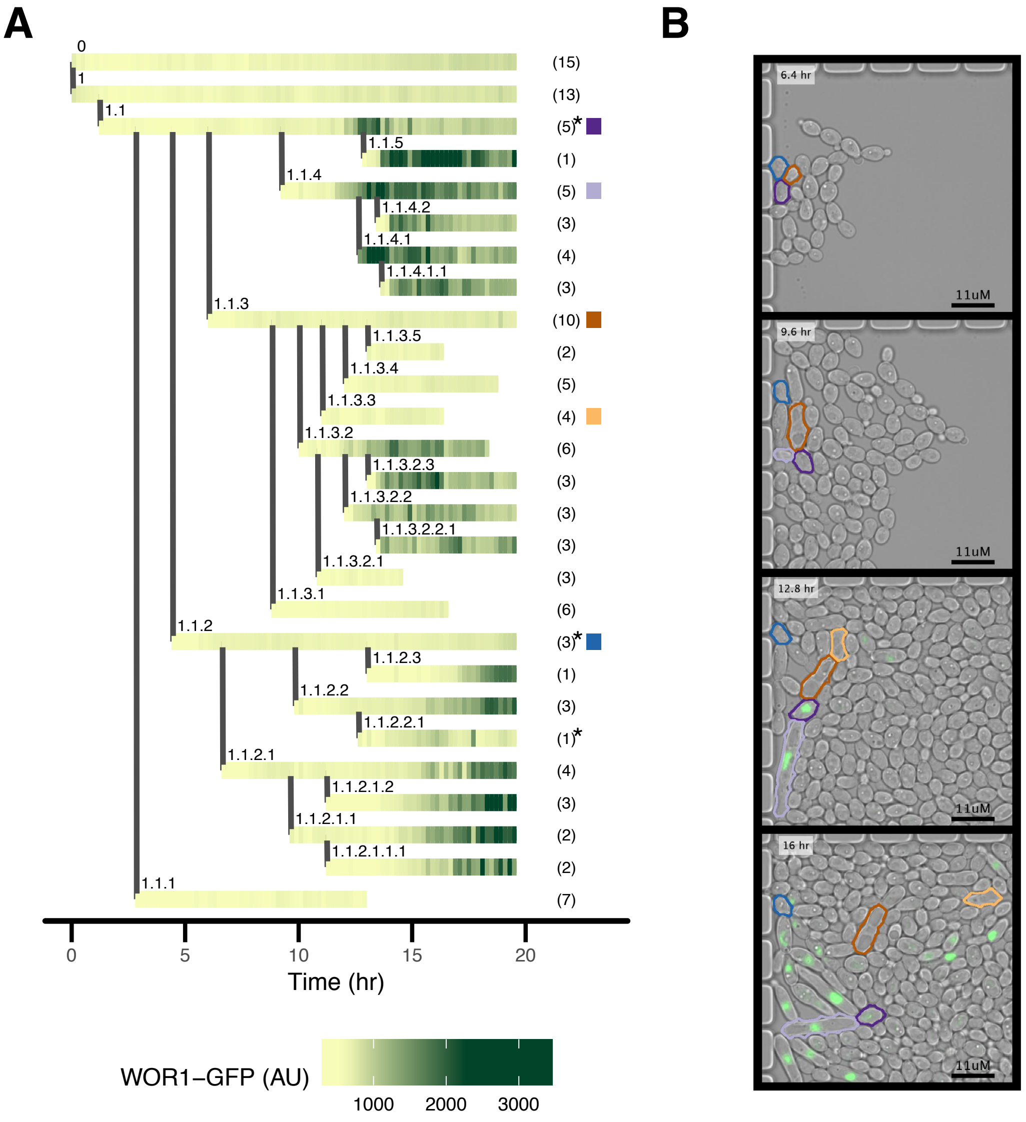


Figure S5 – Mixed fate pedigree example, Wor1-GFP
A) Representative pedigree in which multiple groups of cells activated Wor1. Horizontal lines represent single cells; vertical lines represent budding of a daughter cell. Every horizontal line is made up of small tiles colored by Wor1-GFP fluorescence. Numbers within parentheses on the right of the pedigree represent the number of budded daughter cells per cell within the time period shown; not all daughters are depicted in the pedigree. An asterisk represents that the cell lost its nucleus to its daughter cell. Colored square tiles single out particular cells that are depicted in (B). B) Subset of images representing data shown in (A). Particular cells are outlined in colors corresponding to (A). Cell outlines are based on automated image analysis.

### Figure S6

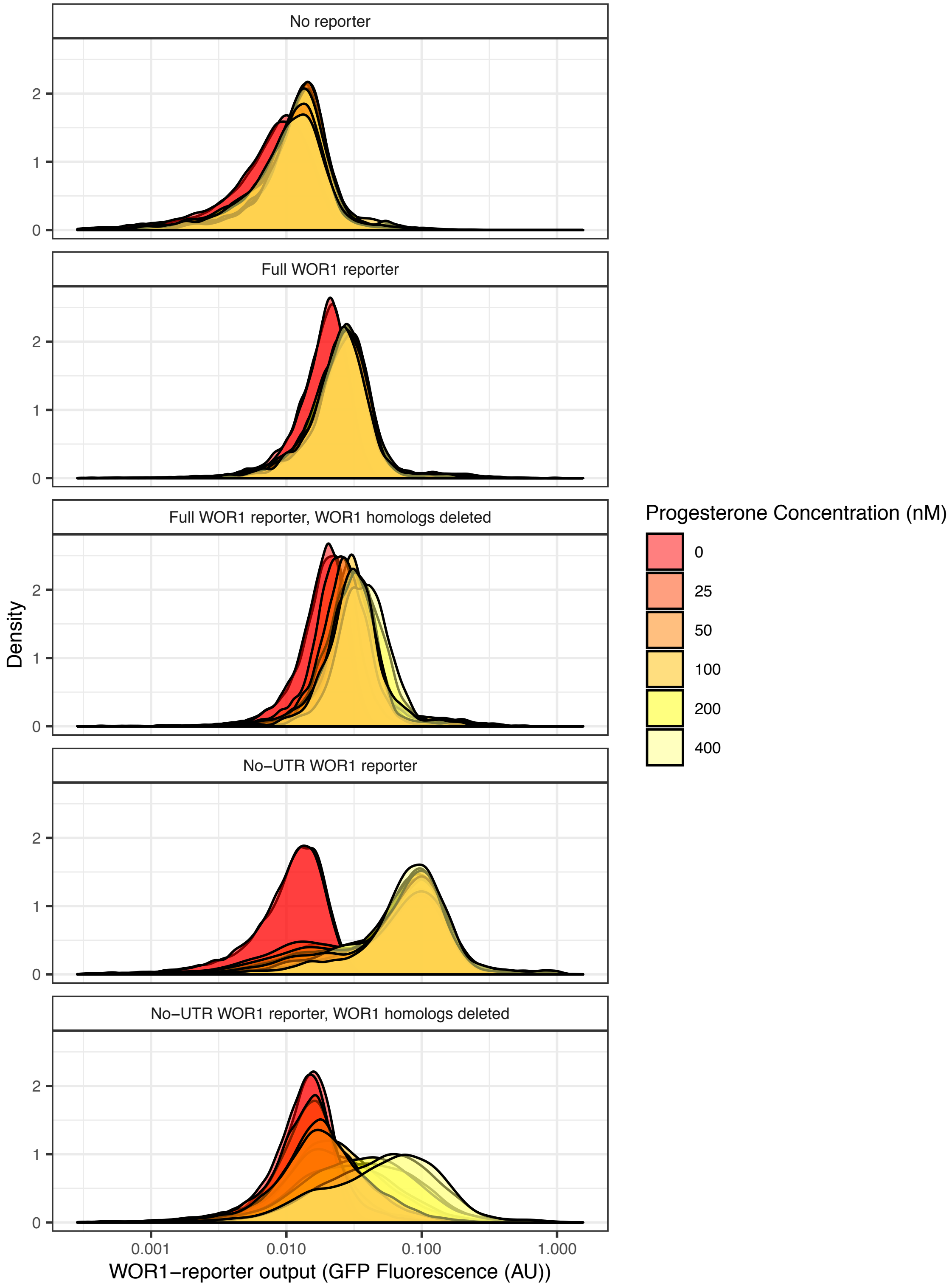


Figure S6 - Alternative Wor1 transcriptional reporters in S. cerevisiae
Distributions of Wor1 transcriptional reporters; all strains shown are inducing Wor1 protein with increasing progesterone concentrations and were independently constructed from the strains shown in the main text. "Full" reporter contains both the Wor1 7KB control region and promoter and the Wor1 2KB 5'UTR. The "No-UTR" reporter contains the Wor1 control region and the Cyc1 core promoter and 5'UTR as explained in the methods section.

### Figure S7

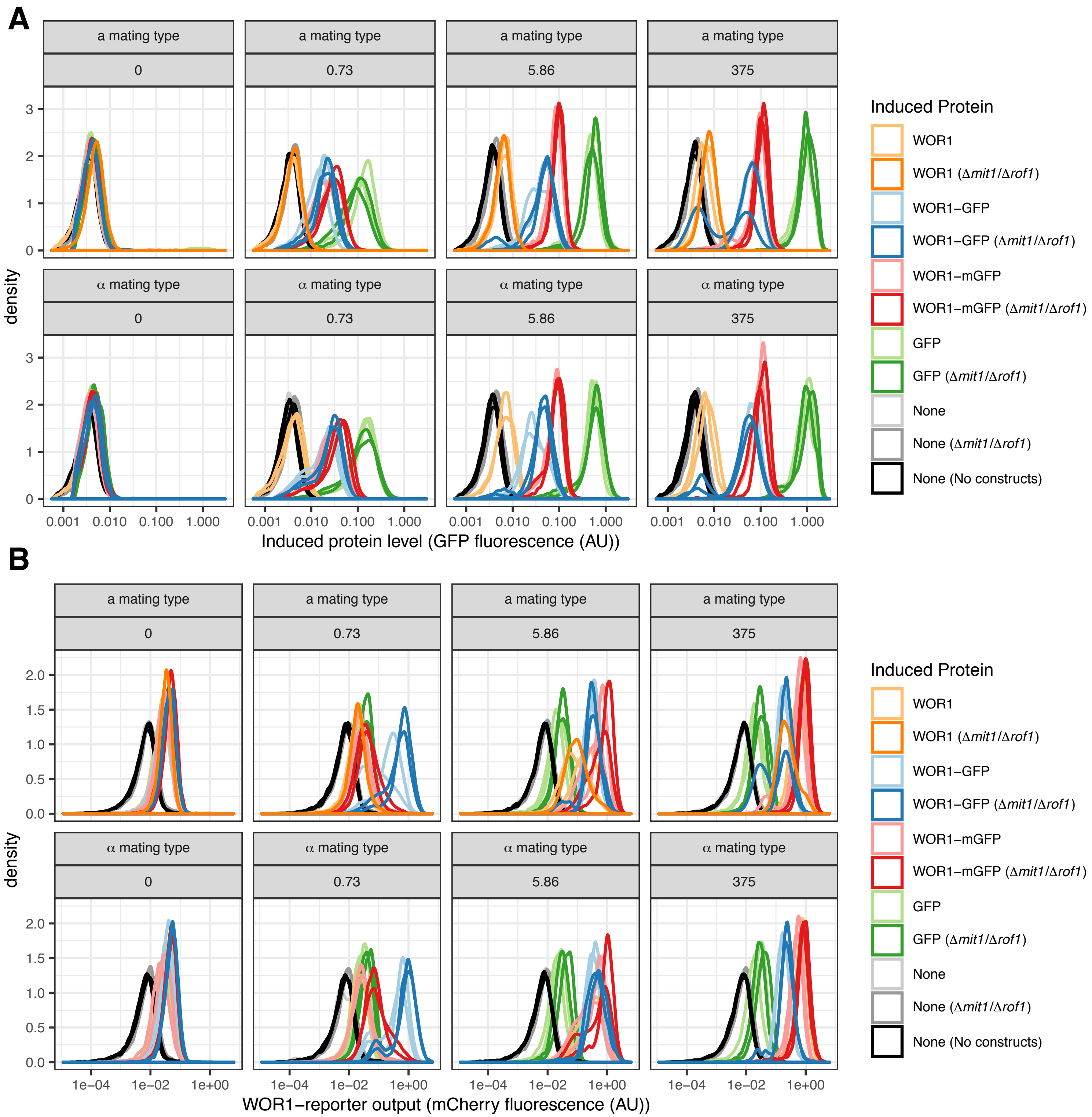


Figure S7 - Distributions of Wor1 protein and reporter levels
Distributions of GFP (A) and mCherry (B) fluorescence; data is shown for all strains in a subset of 4 hormone concentrations (nM progesterone). Note bimodal distributions of GFP fluorescence when inducing Wor1-GFP at high hormone concentrations in (A). Note low-level expression of the Wor1 transcriptional reporter compared to auto-fluorescence in (B).

### Figure S8

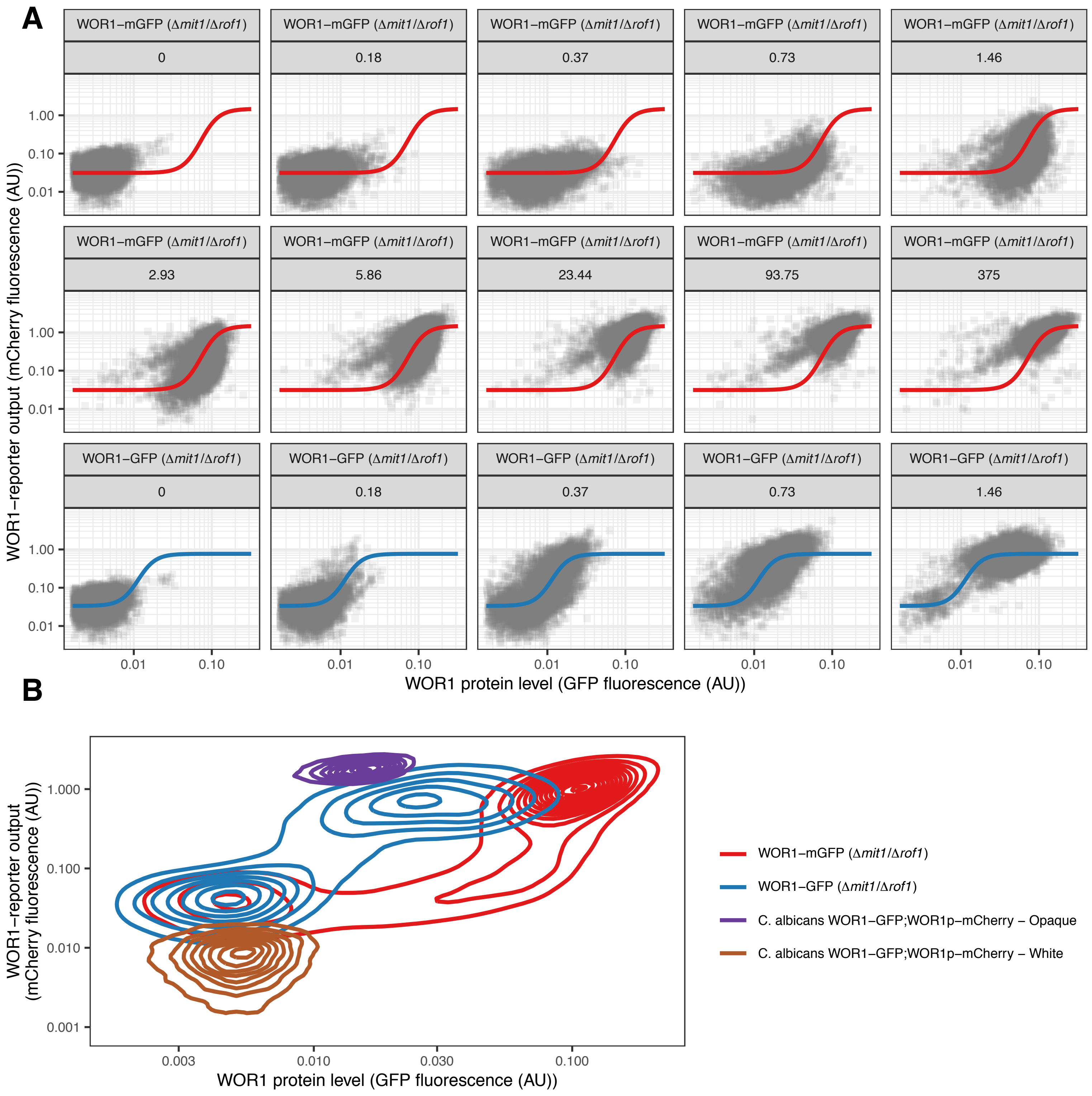


Figure S8 – Distributions of Wor1 protein and reporter levels
A) Same data as Figure 5C, separated by hormone concentration (nM progesterone). B) 2D density plots of flow cytometry data. S. cerevisiae Wor1-GFP and Wor1-mGFP data (same as A) are shown alongside data for white and opaque C. albicans cells containing a Wor1-GFP fusion protein and a transcriptional reporter encompassing the 7KB control region of Wor1 followed by mCherry (no 5'UTR sequence).
